# Genome assembly of the edible jelly fungus *Dacryopinax spathularia* (Dacrymycetaceae)

**DOI:** 10.1101/2024.01.16.575489

**Authors:** Hong Kong Biodiversity Genomics Consortium, Project Coordinator and Co-Principal Investigators, Jerome H.L. Hui, Ting Fung Chan, Leo L. Chan, Siu Gin Cheung, Chi Chiu Cheang, James K.H. Fang, Juan Diego Gaitan-Espitia, Stanley C.K. Lau, Yik Hei Sung, Chris K.C. Wong, Kevin Y.L. Yip, Yingying Wei, DNA extraction, library preparation and sequencing, Tze Kiu Chong, Sean T.S. Law, Genome assembly and gene model prediction, Wenyan Nong, Genome analysis and quality control, Wenyan Nong, Sample collector and logistics, Tze Kiu Chong, Sean T.S. Law, Ho Yin Yip

## Abstract

The edible jelly fungus *Dacryopinax spathularia* (Dacrymycetaceae) is wood-decaying and can be commonly found worldwide. It has also been used in food additives given its ability to synthesize long-chain glycolipids. In this study, we present the genome assembly of *D. spathularia* using a combination of PacBio HiFi reads and Omni-C data. The genome size of *D. spathularia* is 29.2 Mb and in high sequence contiguity and completeness, including scaffold N50 of 1.925 Mb and 92.0% BUSCO score, respectively. A total of 11,510 protein-coding genes, and 474.7 kb repeats accounting for 1.62% of the genome, were also predicted. The *D. spathularia* genome assembly generated in this study provides a valuable resource for understanding their ecology such as wood decaying capability, evolutionary relationships with other fungus, as well as their unique biology and applications in the food industry.

## Introduction

*Dacryopinax spathularia* (Dacrymycetaceae) (Figure 1A) is a brown-rot fungus commonly found on rotting coniferous and broadleaf wood around the world; and can be easily distinguished by the spathulate shape of its gelatinous fruiting body (McNabb, 1965; Worrall et al., 1997). Owing to its production of carotenoid pigments for protection against photodynamic injury, its external appearance is generally orange to yellow (Vail & Lily, 1968). In addition to its ecological role in nutrients recycling, this species is also edible and commonly known as “sweet osmanthus ear” mushroom in China (Bitzer et al., 2018). Given its ability to synthesise long-chain glycolipids under fermentation, this species has also been cultivated in food industry as natural preservatives in soft drinks (EFSA Panel on Food Additives and Flavourings (FAF) et al., 2021).

**Figure 1.**
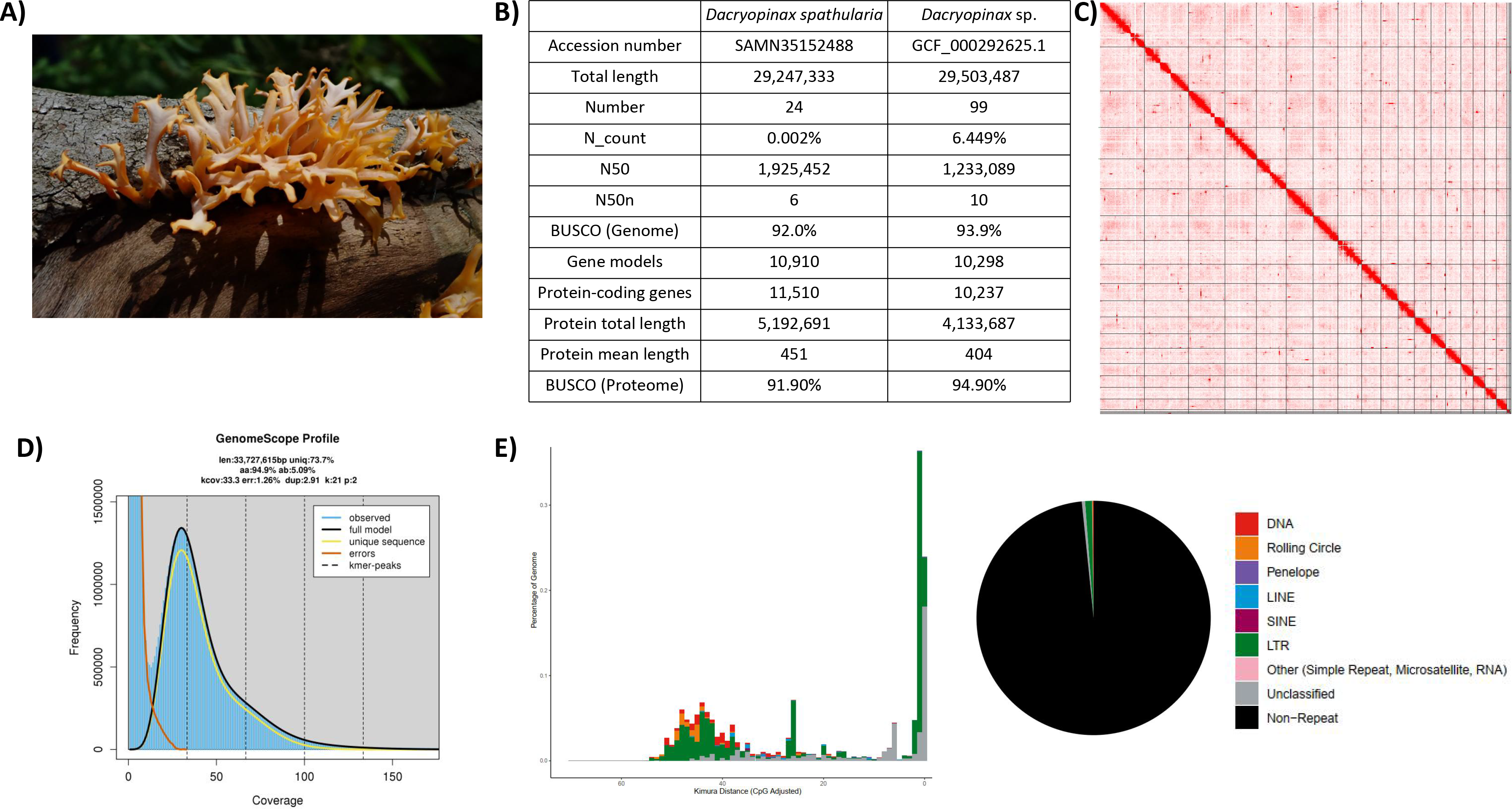
Genomic information of *Dacryopinax spathularia*. **A)** Picture of *Dacryopinax spathularia*; **B)** Genome statistics; **C)** Omni-C contact map of the assembly; **D)** GenomeScope report summary; **E)** Pie chart and repeat landscape plot of repetitive elements in the assembled genome.

## Context

Edible jelly fungus *Dacryopinax spathularia* (Dacrymycetaceae), which was first described as *Merulius spathularius*, is a macrofungus basidiomycete and can be commonly found on rotting coniferous and broadleaf wood in tropics and subtropics. Its wood-decaying ability facilitates nutrient recycling in forest ecosystem (Seifert, 1983). This species is edible and frequently cultivated in industry as food additive such as natural preservative in soft drinks (Bitzer et al., 2018; Bitzer et al., 2019). In addition, isolated fungal extract can also display anti-bacterial properties (Hyde et al., 2019). *D. spathularia* can be naturally found in Asia, Africa, America, Australia and Pacific, and the genomic resource of this species with translational values is not available.

In Hong Kong, *D. spathularia* can also be commonly found (Agriculture, Fisheries and Conservation Department, 2013) and has been selected as one of the species to be sequenced by the Hong Kong Biodiversity Genomics Consortium (a.k.a. EarthBioGenome Project Hong Kong) formed by investigators from eight publicly funded universities. Here, we present the genome assembly of *D. spathularia*, which was assembled from PacBio long reads and Omni-C sequencing data. The provision of the *D. spathularia* genome resource is useful for the better understanding of wood decaying capability, phylogenetic relationships in this family, as well as the biosynthesis of the long-chain glycolipids that are applied as natural preservatives in food industry.

## Methods

### Sample collection and culture of fungal isolates

The fruit bodies of *D. spathularia* was collected in Luk Keng, Hong Kong on 20 June, 2022. The fungal isolate was transferred from the edge of fruit bodies to potato dextrose agar (BD Difco™) plates using a pair of sterilized forceps. The remaining collected fruit bodies were snap-frozen with liquid nitrogen and stored in -80°C refrigerator. Grown hyphae from>2-week-old was then transferred to new plates for purification for at least three rounds. The identity of isolate, termed “F14”, was validated with the DNA barcode of Translation elongation factor 1 alpha (TEF-1α) gene using primer pairs EF1-1018F and EF1-1620R (Stielow et al., 2015) (Supplementary Information 1).

### High molecular weight DNA extraction

∼1.5 g of mycelia of *D. spathularia* isolate was collected from the upper layer of agar culture and was ground in a mortar with liquid nitrogen. High molecular weight (HMW) genomic DNA was isolated with a CTAB treatment, followed by using NucleoBond HMW DNA kit (Macherey Nagel Item No. 740160.20). Briefly, the ground tissue was transferred to 5 mL CTAB buffer (Doyle & Doyle, 1987) with an addition of 1% PVP for 1 h digestion at 55°C. After RNAse A treatment, 1.6 mL 3M potassium acetate was added to the lysate, followed by two rounds of chloroform:IAA (24:1) wash. The supernatant was added with H1 buffer from NucleoBond HMW DNA kit for a final volume of 6 mL and processed according to the manufacturer’s protocol. The DNA sample was eluted in 80 µL elution buffer (PacBio Ref. No. 101-633-500) and its quantity and quality was assessed with NanoDrop^™^ One/OneC Microvolume UV-Vis Spectrophotometer, Qubit^®^ Fluorometer, and overnight pulse-field gel electrophoresis.

### Pacbio library preparation and sequencing

DNA shearing was first performed from 5 µg HMW DNA in 120 µL elution buffer using a g-tube (Covaris Part No. 520079) with 6 passes of centrifugation at 1,990 x *g* for 2 min. The sheared DNA sample was purified using SMRTbell^®^ cleanup beads (PacBio Ref. No. 102158-300), from which 2 µL of sample was taken for quality check through overnight pulse-field gel electrophoresis and Qubit^®^ Fluorometer quantification. A SMRTbell library was then prepared by following the protocol of the SMRTbell® prep kit 3.0 (PacBio Ref. No. 102-141-700). Briefly, the sheared DNA was repaired and polished at both ends, followed by A-tailing and ligation of T-overhand SMRTbell adapters. A subsequent purification step was processed with SMRTbell^®^ cleanup beads and 2 µL of sample was taken and subject to quality check as mentioned above. Nuclease treatment was then proceeded to remove non-SMRT bell structures. A final size-selection step using 35% AMPure PB beads was processed to eliminate short fragments.

A final library preparation was performed with The Sequel^®^ II binding kit 3.2 (PacBio Ref. No. 102-194-100) before sequencing. The SMRTbell library was proceeded with annealing and binding with Sequel II^®^ primer 3.2 and Sequel II^®^ DNA polymerase 2.2, respectively. SMRTbell^®^ cleanup beads were used for further cleanup the library, to which diluted Sequel II^®^ DNA Internal Control Complex was added. The final library was loaded at an on-plate concentration of 90 pM with the diffusion loading mode. Sequencing was performed on the Pacific Biosciences SEQUEL IIe System for a run of 30-hour movies with 120 min pre-extension to output HiFi reads with one SMRT cell. Details of the resulting sequencing data are listed in Supplementary Information 2.

### Omnic-C library preparation and sequencing

∼0.5 g of stored fruit body was ground into powder with liquid nitrogen and used for the construction of an Omni-C library by following the plant tissue protocol of the Dovetail® Omni-C® Library Preparation Kit (Dovetail Cat. No. 21005). The ground tissue was transferred to 4 mL 1X PBS and was proceeded to crosslinking with formaldehyde and digestion with endonuclease DNase I. The quantity and fragment size of the lysate was assessed with Qubit^®^ Fluorometer and TapeStation D5000 HS ScreenTape, respectively. The qualified lysate was polished at DNA ends and ligated with biotinylated bridge adaptors, followed by proximity ligation, crosslink reversal of DNA and purification with SPRIselect™ Beads (Beckman Coulter Product No. B23317). The end repair and adapter ligation were performed with the Dovetail™ Library Module for Illumina (Dovetail Cat. No. 21004). The library was then sheared with USER Enzyme Mix and purified with SPRIselect™ Beads. The DNA fragments were isolated in Streptavidin Beads, from which the library was amplified with Universal and Index PCR Primers from the Dovetail™ Primer Set for Illumina (Dovetail Cat. No. 25005). Size selection targeting fragment size between 350 bp and 1000 bp was performed with SPRIselect™ Beads. The quantity and fragment size of the library was assessed by Qubit^®^ Fluorometer and TapeStation D5000 HS ScreenTape, respectively. The resulting library was sequenced on an Illumina HiSeq-PE150 platform. Details of the resulting sequencing data are listed in Supplementary Information 2.

### RNA extraction and transcriptome sequencing

∼1 g of mycelia of *D. spathularia* isolate was ground in a mortar with liquid nitrogen. Total RNA was isolated from the ground tissue using the mirVana miRNA Isolation Kit (Ambion), following the manufacturer’s instructions. The RNA sample was subjected to quality control with NanoDrop^™^ One/OneC Microvolume UV-Vis Spectrophotometer and 1% agarose gel electrophoresis. The qualified sample was sent to Novogene Co. Ltd (Hong Kong, China) for 150 bp paired-end sequencing. Details of the resulting sequencing data are listed in Supplementary Information 2.

### Genome assembly and gene model prediction

A *de novo* genome assembly was conducted with Hifiasm (Cheng et al., 2021), which was screened with BlobTools (v1.1.1) (Laetsch & Blaxter, 2017) by searching against the NT database using BLAST to identify and remove any possible contaminations (Supplementary Information 3). Haplotypic duplications were discarded using “purge_dups” according to the depth of HiFi reads (Guan et al., 2020). The Omni-C data were used to scaffold the assembly using YaHS (Zhou et al., 2022).

Gene model prediction was performed using funannotate (Palmer & Stajich, 2020). RNA sequencing data were first processed using Trimmomatic (v0.39) and kraken2 (v2.0.8 with kraken2 database k2_standard_20210517) to remove the low quality and contaminated reads. The processed reads were then aligned to the soft-masked repeat genome using Hisat2 to run the genome-guided Trinity (Grabherr et al., 2011) with parameters ”--stranded RF--jaccard_clip”, from which 44,384 transcripts were derived. Gene models were then predicted together with the protein evidence from *Dacryopinax primogenitus* (GCF_000292625.1; Floudas et al., 2012) using funannotate with the following parameters “--protein_evidenceGCF_000292625.1_Dacryopinax_sp._DJM_731_SSP1_v1.0.proteins.faa--genemark_mode ET --optimize_augustus --busco_db dikarya --organism fungus -d--max_intronlen 3000”. The Trinity transcript alignments were converted to GFF3 format and were input to PASA alignment in the Launch_PASA_pipeline.pl process to generate the PASA models trained by TransDecoder, followed by selection of the PASA gene models using the Kallisto TPM data. The PASA gene models were then used for training Augustus in the funannotate-predict step. The gene models from several prediction sources, with a total of 54,275 genes from Augustus (4967), HiQ (4624), CodingQuarry (11762), GlimmerHMM (10843), pasa (11217), snap (10862), were passed to Evidence Modeler to generate the gene model annotation files. UTRs were then captured in the funannotate-update step using PASA to generate the final genome annotation files.

### Repeat annotation

Transposable element (TE) annotation was performed by following the Earl Grey TE annotation workflow pipeline (version 1.2, https://github.com/TobyBaril/EarlGrey) (Baril et al., 2022).

## Results and discussion

### Genome assembly

A total of 9.34 Gb HiFi reads were generated from PacBio sequencing (Supplementary Information 2). After scaffolding with 3.46 Gb Omni-C data, the *D. spathularia* genome assembly has a size of 29.2 Mb, scaffold N50 of 1.925 Mb and 92.0% BUSCO score (Figure 1B; Table 1), and 19 out of 24 scaffolds are >100 kb in length (Figure and 1C; Table 2). The genome size is similar to *Dacryopinax primogenitus* (29.5 Mb) (Floudas et al., 2012) and GenomeScope estimated heterozygosity of 5.09% (Figure 1D; Table 3). Gene model prediction generated a total of 11,510 protein-coding genes with an average protein length of 451 bp and a BUSCO score of 91.9%.

### Repeat content

Repeat content analysis showed that transposable elements (TEs) account for 1.62% of the *D. spathularia* genome (Figure 1E; Table 4). The major classified TE was LTR retransposons (0.95%) and DNA transposons (0.12%) (Table 4).

### Conclusion and future perspective

The study presents the genome assembly of *D. spathularia*, which is a useful resource for further phylogenomic studies in the family Dacrymycetaceae and investigations on the biosynthesis of glycolipids via the fermentation process with applications in the food industry.

### Data validation and quality control

The identity of fungal isolate of *D. spathularia* was validated with DNA barcoding of Translation elongation factor 1 alpha (TEF-1α) gene, which was compared with sequences from phylogenetic studies of Dacrymycetaceae (Zaroma & Ekman, 2020) and its sister family Cerinomycetaceae (Savchenko et al., 2021), the *Dacryopinax primogenitus* genome (Accession: NW_024467206.1:736197-736766), and *D. spathularia* (Accession: AY881020.1). The sequences were aligned with MAFFT v7.271 (Katoh and Standley, 2013). A phylogenetic tree was constructed with FastTree (Price et al., 2010) with 1,000 bootstraps and visualized in Evolview v3 (Subramanian et al., 2019). The *D. spathularia* isolate in this study was clustered with other two *D. spathularia* accessions with a bootstrap support of 92/100 (Supplementary Information 1).

For HMW DNA extraction and Pacbio library preparation, the samples were subject to quality control with NanoDrop^™^ One/OneC Microvolume UV-Vis Spectrophotometer, Qubit^®^ Fluorometer, and overnight pulse-field gel electrophoresis. The quality of Omni-C library was inspected with Qubit^®^ Fluorometer and TapeStation D5000 HS ScreenTape.

During the genome assembly, BlobTools (v1.1.1) (Laetsch & Blaxter, 2017) was employed to identify and remove any possible contaminations (Supplementary Information 3). The assembled genome and gene model prediction were assessed with Benchmarking Universal Single-Copy Orthologs (BUSCO, v5.5.0) (Manni et al., 2021) using the fungi dataset (fungi_odb10). GenomeScope2 (Vurture et al., 2017) was used to estimate the genome size and heterozygosity of the assembly.

## Data availability

The raw reads generated in this study were deposited in the NCBI database under the SRA accessions SRR24631918, SRR27412332 and SRR27412333. The GenomeScope report, genome, genome annotation and repeat annotation files were made publicly available in Figshare (https://figshare.com/s/9d7dd8509b902306bd5b).

## Authors’ contribution

JHLH, TFC, LLC, SGC, CCC, JKHF, JDG, SCKL, YHS, CKCW, KYLY and YW conceived and supervised the study; TKC and STSL collected the samples and carried out DNA extraction, library preparation and genome sequencing; HYY arranged the logistics of samples; WN performed genome assembly and gene model prediction.

---

**Table 1**. Genome statistic and sequencing information.

**Table 2**. Information on scaffold name and length.

**Table 3**. Summary of GenomeScope statistics.

**Table 4**. Summary of transposable element annotation.

**Supplementary Information 1**. Phylogenetic analysis of Translation elongation factor 1 alpha (TEF-1α) gene region *Dacryopinax spathularia* fungal isolate “F14” in this study.

**Supplementary Information 2**. Summary of genomic sequencing data.

**Supplementary Information 3**. Genome assembly QC and contaminant/cobiont detection.

## References

1. Agriculture, Fisheries and Conservation Department. Common wood decay fungi of Hong Kong (2). 2013. https://www.herbarium.gov.hk/filemanager/leaflets/en/upload/3/13/7_en.pdf. Accessed 04 Jan 2024.

2. Baril T, Imrie RM, Hayward A. Earl Grey: a fully automated user-friendly transposable element annotation and analysis pipeline. bioRxiv. 2022.10.1101/2022.06.30.498289

3. Bitzer J, Henkel T, Nikiforov AI, Rihner MO, Herberth MT. Developmental and reproduction toxicity studies of glycolipids from Dacryopinax spathularia. Food and Chemical Toxicology. 2018;120:430–8.

4. Bitzer J, Henkel T, Nikiforov AI, Rihner MO, Verspeek-Rip CM, Usta B, van den Wijngaard M. Genetic toxicity studies of glycolipids from Dacryopinax spathularia. Food and Chemical Toxicology. 2019;123:162–8.

5. Cheng H, Concepcion GT, Feng X, Zhang H, Li H. Haplotype-resolved de novo assembly using phased assembly graphs with hifiasm. Nature methods. 2021;18(2):170–5.

6. Doyle JJ, Doyle JL. A rapid DNA isolation procedure for small quantities of fresh leaf tissue. Phytochemical bulletin. 1987.

7. EFSA Panel on Food Additives and Flavourings (FAF), Younes M, Aquilina G, Engel KH, Fowler P, Frutos Fernandez MJ, Fürst P, Gürtler R, Gundert[Remy U, Husøy T, Manco M. Safety evaluation of long chain glycolipids from Dacryopinax spathularia. EFSA Journal. 2021;19(6):e06609.

8. Floudas D, Binder M, Riley R, Barry K, Blanchette RA, Henrissat B, Martínez AT, Otillar R, Spatafora JW, Yadav JS, Aerts A. The Paleozoic origin of enzymatic lignin decomposition reconstructed from 31 fungal genomes. Science. 2012;336(6089):1715–9.

9. Guan D, McCarthy SA, Wood J, Howe K, Wang Y, Durbin R. Identifying and removing haplotypic duplication in primary genome assemblies. Bioinformatics. 2020;36(9):2896–8.

10. Grabherr MG, Haas BJ, Yassour M, Levin JZ, Thompson DA, Amit I, Adiconis X, Fan L, Raychowdhury R, Zeng Q, Chen Z. Full-length transcriptome assembly from RNA-Seq data without a reference genome. Nature biotechnology. 2011;29(7):644–52.

11. Hyde KD, Xu J, Rapior S, Jeewon R, Lumyong S, Niego AG, Abeywickrama PD, Aluthmuhandiram JV, Brahamanage RS, Brooks S, Chaiyasen A. The amazing potential of fungi: 50 ways we can exploit fungi industrially. Fungal Diversity. 2019;97:1–36.

12. Katoh K, Standley DM. MAFFT multiple sequence alignment software version 7: improvements in performance and usability. Molecular biology and evolution. 2013;30(4):772–80.

13. Laetsch DR, Blaxter ML. BlobTools: Interrogation of genome assemblies. F1000Research. 2017;6(1287):1287.

14. Manni M, Berkeley MR, Seppey M, Simão FA, Zdobnov EM. BUSCO update: novel and streamlined workflows along with broader and deeper phylogenetic coverage for scoring of eukaryotic, prokaryotic, and viral genomes. Molecular biology and evolution. 2021;38(10):4647–54.

15. McNabb RF. Taxonomic studies in the Dacrymycetaceae: III. Dacryopinax Martin. New Zealand Journal of Botany. 1965;3(1):59–72.

16. Nagy LG, Riley R, Tritt A, Adam C, Daum C, Floudas D, Sun H, Yadav JS, Pangilinan J, Larsson KH, Matsuura K. Comparative genomics of early-diverging mushroom-forming fungi provides insights into the origins of lignocellulose decay capabilities. Molecular biology and evolution. 2016;33(4):959–70.

17. Palmer JM, Stajich J. Funannotate v1. 8.1: Eukaryotic genome annotation. Zenodo 10.5281/zenodo.2020;4054262.

18. Price MN, Dehal PS, Arkin AP. FastTree 2–approximately maximum-likelihood trees for large alignments. PloS one. 2010;5(3):e9490.

19. Savchenko A, Zamora JC, Shirouzu T, Spirin V, Malysheva V, Kõljalg U, Miettinen O. Revision of Cerinomyces (Dacrymycetes, Basidiomycota) with notes on morphologically and historically related taxa. Studies in Mycology. 2021;99(1):100117.

20. Seifert KA. Decay of wood by the Dacrymycetales. Mycologia. 1983:1011–8.

21. Shirouzu T, Hosaka K, Nam KO, Weir BS, Johnston PR, Hosoya T. Phylogenetic relationships of eight new Dacrymycetes collected from New Zealand. Persoonia-Molecular Phylogeny and Evolution of Fungi. 2017;38(1):156–69.

22. Stielow JB, Levesque CA, Seifert KA, Meyer W, Irinyi L, Smits D, Renfurm RG, Verkley GJ, Groenewald M, Chaduli D, Lomascolo A. One fungus, which genes? Development and assessment of universal primers for potential secondary fungal DNA barcodes. Persoonia-Molecular Phylogeny and Evolution of Fungi. 2015;35(1):242–63.

23. Subramanian B, Gao S, Lercher MJ, Hu S, Chen WH. Evolview v3: a webserver for visualization, annotation, and management of phylogenetic trees. Nucleic acids research. 2019;47(W1):W270–5.

24. Vail WJ, Lilly VG. The location of carotenoid pigments and thickness of the cell wall in light-and dark-grown cells of Dacryopinax spathularia. Mycologia. 1968;60(4):902–7.

25. Vurture GW, Sedlazeck FJ, Nattestad M, Underwood CJ, Fang H, Gurtowski J, Schatz MC. GenomeScope: fast reference-free genome profiling from short reads. Bioinformatics. 2017;33(14):2202–4.

26. Worrall JJ, Anagnost SE, Zabel RA. Comparison of wood decay among diverse lignicolous fungi. Mycologia. 1997;89(2):199–219.

27. Zamora JC, Ekman S. Phylogeny and character evolution in the Dacrymycetes, and systematics of Unilacrymaceae and Dacryonaemataceae fam. nov. Persoonia-Molecular Phylogeny and Evolution of Fungi. 2020;44(1):161–205.

28. Zhou C, McCarthy SA, Durbin R. YaHS: yet another Hi-C scaffolding tool. Bioinformatics. 2023;39(1):btac808.

