## Supplementary material for "Genome assembly of the edible jelly fungus *Dacryopinax spathularia* (Dacrymycetaceae)": EPBHK_Fungi Supplementary Information.docx

**Supplementary Information 1.** Phylogenetic analysis of Translation elongation factor 1 alpha (TEF-1α) gene region *Dacryopinax spathularia* fungal isolate “F14” in this study. Sequences from related phylogenetic studies and NCBI accessions were incorporated, including Zaroma & Ekman (2020), Savchenko et al. (2021), *Dacryopinax primogenitus* genome (NW_024467206.1:736197-736766), and NCBI BLAST result of a *D. spathularia* isolate (Accession: AY881020.1), which are highlighted in light blue, pink, green, and red, respectively. The fungal isolate “F14” in this study is highlighted in orange. The bootstrap values in percentage are shown at the nodes.

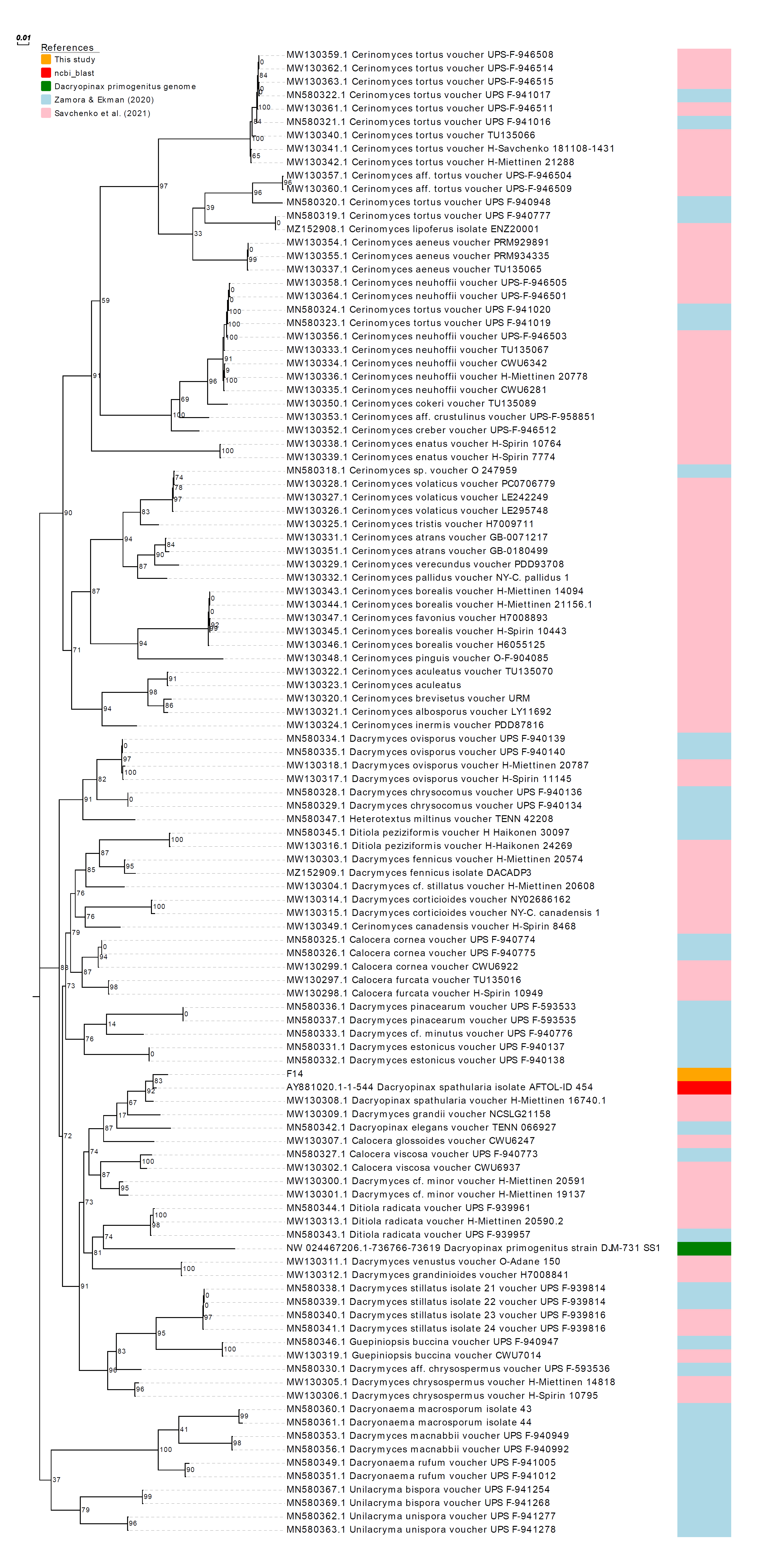

Cerinomycetaceae

Dacrymycetaceae

Unilacrymaceae

**Supplementary Information 2.** Genome and transcriptome sequencing data.

| Library | Reads | Bases | Coverage (X) | Accession number |
| --- | --- | --- | --- | --- |
| PacBio HiFi | 757,219 | 9,340,525,111 | 319 | SRR24631918 |
| Omnic | 23,101,878 | 3,465,281,700 | 118 | SRR27412332 |
| mRNA | 39,483,524 | 5,922,432,049 | 202 | SRR27412333 |

**Supplementary Information 3.** Genome assembly QC and contaminant/cobiont detection.

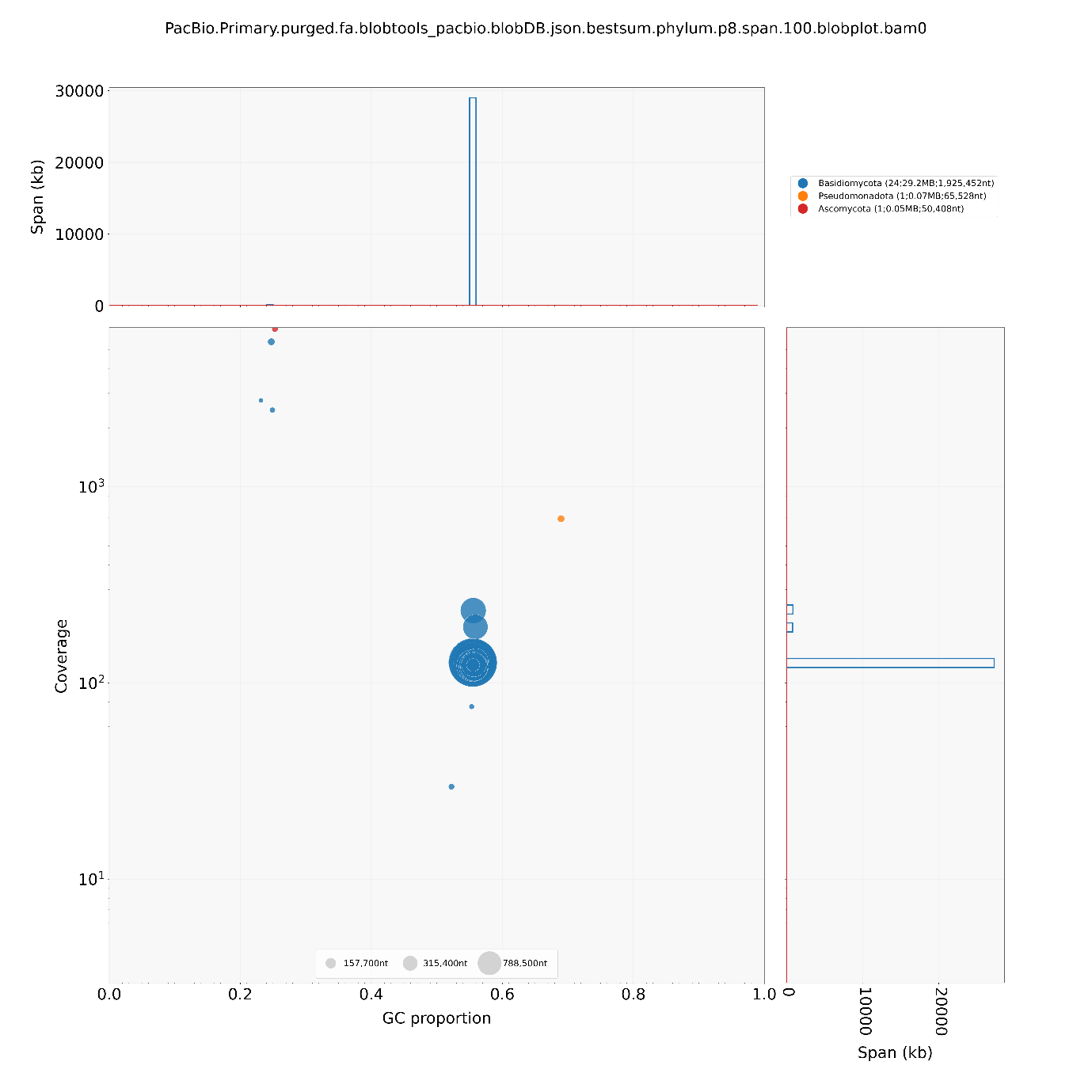

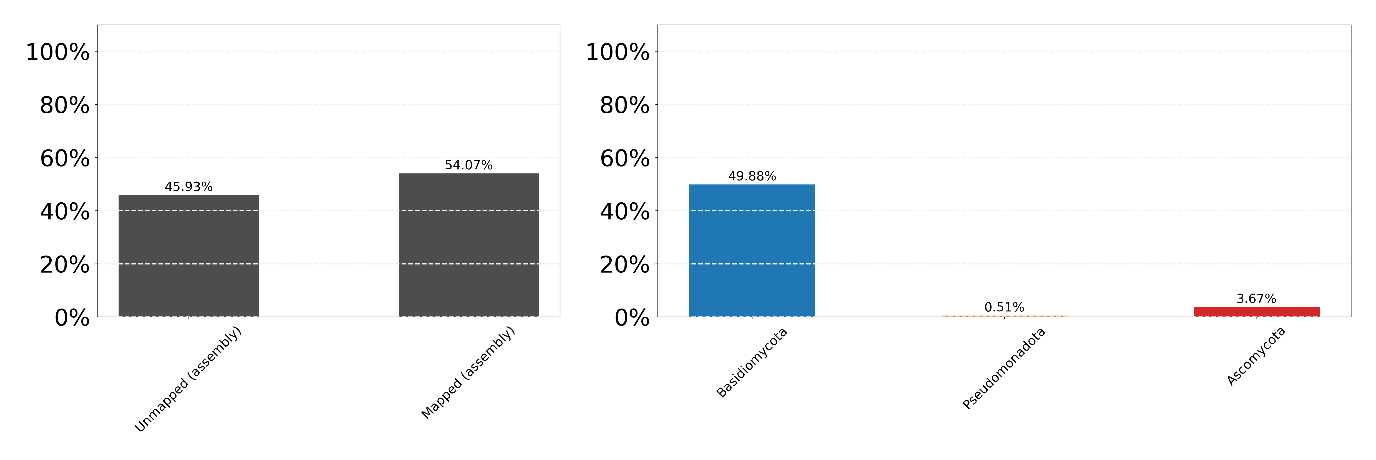
